# RNA virome diversity and *Wolbachia* infection in individual *Drosophila simulans* flies

**DOI:** 10.1101/2021.05.09.443333

**Authors:** Ayda Susana Ortiz-Baez, Mang Shi, Ary A. Hoffmann, Edward C. Holmes

## Abstract

The endosymbiont bacterium *Wolbachia* is associated with multiple mutualistic effects on insect biology, including nutritional and antiviral properties. *Wolbachia* naturally occurs in *Drosophila* fly species, providing an operational model host to study how virome composition may be impacted by its presence. *Drosophila simulans* populations can carry a variety of *Wolbachia* strains. In particular, the *w*Au strain of *Wolbachia* has been associated with strong antiviral protection under experimental conditions. We used *D. simulans* sampled from the Perth Hills, Western Australia, to investigate the potential virus protective effect of the *w*Au strain on individual wild-caught flies. Our data revealed no appreciable variation in virus composition and abundance between *Wolbachia* infected/uninfected individuals associated with the presence/absence of *w*Au. However, it remains unclear whether *w*Au might impact viral infection and host survival by increasing tolerance rather than inducing complete resistance. These data also provide new insights into the natural virome diversity of *D. simulans*. Despite the small number of individuals sampled, we identified a repertoire of RNA viruses, including Nora virus, Galbut virus, Chaq virus, Thika virus and La Jolla virus, that have been identified in other *Drosophila* species. In addition, we identified five novel viruses from the families *Reoviridae*, *Tombusviridae*, *Mitoviridae* and *Bunyaviridae.* Overall, this study highlights the complex interaction between *Wolbachia* and RNA virus infections and provides a baseline description of the natural virome of *D. simulans*.

## Introduction

The alpha-proteobacterium *Wolbachia* (order *Rickettsiales*) is a widespread endosymbiont of arthropods and nematodes (i.e. filarial and plant-parasitic nematodes), that can establish interactions with their hosts ranging from parasitic to mutualistic [1,2]. The genetic diversity of *Wolbachia* is substantial and currently represented by 11 distinctive supergroups (denoted A-J), although the majority of *Wolbachia* strains belong to supergroups A and B [3] that are estimated to have diverged around 50 million years ago [4]. Although these bacteria are commonly found in reproductive tissues and the germline of their hosts, they have also been found in somatic tissues such as the brain, salivary glands and gut [5–9], such that understanding infection dynamics in detail is not a trivial matter [7]. *Wolbachia* primarily spread by vertical inheritance through transovarian transmission. However, the presence of *Wolbachia* in a diverse range of host species suggests that horizontal transmission, likely through antagonistic interactions (i.e. herbivory, parasitism and predation), also contributes to the dissemination of the bacteria in nature [4,10].

The occurrence of *Wolbachia* bacteria in insects is often associated with their ability to manipulate host reproductive mechanisms and induce a range of alterations, including parthenogenesis, feminization, cytoplasmic incompatibility and sex-ratio distortion [11]. Among these, cytoplasmic incompatibility is the most common phenotypic effect, and as such represents an appealing approach for vector population control. In this case, embryonic lethality is contingent on the infection status and the strain type harboured by males and females [2]. In addition, the study of *Wolbachia*-host interactions has revealed a variety of mutualistic effects on host biology [1,12]. For instance, in filarial nematodes and the parasitoid wasp *Asobara tabida*, the presence of some *Wolbachia* strains has been positively associated with developmental processes, fertility and host viability [12–14]. Furthermore, nutritional mutualism between *Wolbachia* and the bedbug *Cimex lectularius* as well as *Wolbachia*-infected planthoppers, has been suggested as a means to explain B vitamin supplementation [15–17].

Arguably the most important outcome of *Wolbachia* infection in insects is its potential for virus-blocking, which also provides a basis for intervention strategies based on the control of arbovirus transmission. This seemingly antiviral effect of *Wolbachia* has been well documented in some species of insects, including flies and mosquitoes. A striking example involves the transinfection of *Aedes aegypti* mosquitoes with the *Wolbachia* strain infecting *Drosophila melanogaster* (*w*Mel)*. A. aegypti* is the primary vector of a number of important arboviruses, including Dengue (DENV), Zika (ZIKV) and Chikungunya virus (CHIKV), and the establishment of the *w*Mel strain in wild mosquito populations represents a powerful and promising approach to decrease virus transmission [18,19]. Although the underlying mechanisms remain to be fully determined, it has been suggested that *Wolbachia* can modify the host environment or boost basal immunity to viruses by pre-stimulating the immune response of their hosts [20]. Potential antiviral mechanisms impacted by *Wolbachia* include gene expression of the Toll pathway, RNA interference, and modification of the host oxidative environment that likely trigger an antiviral immune response and hence limit infection [20–22].

Unlike *A. aegypti* mosquitoes, *Wolbachia* naturally occur in *Drosophila* species, providing a valuable model system to study *Wolbachia*-related virus protection [23,24]. Natural populations of *Drosophila* can carry a diverse array of insect-specific viruses belonging to the families *Picornaviridae*, *Dicistroviridae*, *Bunyaviridae*, *Reoviridae* and *Iflaviridae* amongst others [25]. The co-occurrence of *Wolbachia* in *D. melanogaster* has been associated with increased survival and different levels of resistance to laboratory viral infections in fly stocks under experimental conditions [23,26]. For example, *Wolbachia*-infected flies containing the dicistrovirus *Drosophila C virus* (DCV) showed a delay in mortality compared to *Wolbachia*-free flies [26]. In contrast, other studies found no or limited effect of *Wolbachia* on viral protection, as well as on virus prevalence and abundance in field-collected flies [25,27]. Such contrasting data emphasize the need of further research efforts to characterize the effect of *Wolbachia* strains on virus composition in *Drosophila* in nature.

Although the origin of *D. simulans* is thought to have been in East Africa or Madagascar, this species now has a cosmopolitan distribution [28]. In Australia, *D. simulans* has been recorded along both east and west coasts as well as Tasmania, with the earliest record dating to 1956 [29]. Human mobility and human-mediated activities have been associated with the introduction and spread of both *D. simulans* and *Wolbachia* into Australia, where wild fly populations occur near human settlements, feeding and breeding on a variety horticultural crops [30,31]. Several *Wolbachia* strains from supergroups A and B can naturally occur in populations of *D. simulans* (e.g. *w*Au, *w*Ri, *w*Ha, *w*Ma and *w*No) [32,33]. From these, *w*Au is associated with strong antiviral protection against Flock House virus (*Nodaviridae*) and Drosophila C virus (*Dicistroviridae*) under experimental conditions [32]. The *w*Au infection in Australia was one of the first *Wolbachia* infections identified as showing no cytoplasmic incompatibility despite being widespread at a low to intermediate frequency [34]. *w*Au was subsequently demonstrated to be increasing in frequency along the east coast of Australia, until it was replaced by *w*Ri that shows cytoplasmic incompatibility but which has not yet reached the west coast [30]. In this study, we used a meta-transcriptomic (i.e. RNA shotgun sequencing) approach to determine the virome diversity of individual field-collected *D. simulans* flies from Western Australia, and investigated how this virome diversity might be impacted by the presence of the *w*Au strain of *Wolbachia*.

## Methods

### D. simulans collection and taxonomic identification

Flies used for the virus work were collected at Raeburn Orchards in the Perth Hills in Western Australia (Long. 116.0695, Lat. −32.1036) in July 2018 using banana bait. The *Wolbachia* frequency at two other locations in the area (Roleystone, Long. 116.0701, Lat. −32.1396; Cannington, Long. 115.9363, Lat. −32.0243) was also established with additional samples. Taxonomic identification to the species level was conducted based on the morphology of reproductive traits of males and via DNA barcoding. Field-collected flies were maintained at 19°C under standard laboratory conditions until F1 offspring were raised. Parental and F1 generations were then stored at −80°C until molecular processing.

### Wolbachia detection

*Wolbachia* infection of field females was determined using F1 offspring from each field female. Note that wAu is transmitted at 100% from field females to the F1 laboratory generation [34]. DNA extraction from heads was performed using the Chelex 100 Resin (Bio-Rad Laboratories, Hercules, CA, USA) [35] as adapted in Shi *et al*. [27]. Screening of natural *Wolbachia* infection was conducted using a real-time PCR/ high-resolution melt assay (RT/HRM) and strain-specific primers targeting a 340-bp region of the surface protein of Wolbachia (*wsp)* gene for *w*Ri and *w*Au strains. The assay was run following the protocol of Kriesner *et al*. [30]. In addition, reads were mapped to reference *Wolbachia wsp* gene sequences for *w*Ri (CP001391.1) and *w*Au (LK055284.1) with BBMap v.37.98 (minid=0.95) (available at https://sourceforge.net/projects/bbmap/).

### RNA extraction and meta-transcriptome sequencing

We screened a total of 16 individual flies to assess the effect of *Wolbachia* infection on virome composition in *D. simulans*. Specimens were rinsed three times in RNA and DNA-free PBS solution (GIBCO). Total RNA from individual flies was extracted using the RNeasy Plus Mini Kit (Qiagen) following the manufacturer’s instructions. RNA-seq libraries were constructed using a TruSeq total RNA Library Preparation Kit (Illumina). Host ribosomal depletion was performed using a Ribo-Zero Gold rRNA Removal Kit (Human/Mouse/Rat) (Illumina). Paired-end transcriptome sequencing was generated on the HiSeq2500 platform (Illumina).

### De novo meta-transcriptome assembly and viral genome annotation

The overall quality assessment of reads was conducted in FastQC and Trimmomatic [36]. A *de novo* assembly of RNA-Seq data was performed using MEGAHIT v.1.1.3, with default parameters [37]. Assembled contigs were then annotated through comparisons against the NCBI nonredundant (NCBI-nr) database using DIAMOND v2.0.4 [38], with a cut-off e-value <1e-05. To identify protein-encoding sequences, open reading frames (ORFs) were predicted in positive and reverse-complement strands, with a minimum length of 600 nt between two stop codons using the GetOrf program (EMBOSS) [39]. Functional annotation was carried out using InterProScan v5.39-77.0 [40], and the HMMer software (http://hmmer.org/) was used to perform sequence-profile searches against the Pfam HMM database. To expand the *de novo* assembled contigs of known viruses, the reads were mapped against reference genomic sequences. Provisional virus names were derived from geographic locations in the Perth Hills.

### Estimates of viral abundance

Viral abundance was assessed using the number of reads per million (RPM). This metric quantifies the number of reads per million mapped to a given contig assembly over the total number of reads. RPM values lower than 0.1% of the highest count for each virus across samples were presumed to be index-hopping artifacts and excluded from the remaining analyses [41]. To compare abundance levels, reads were mapped to reference ribosomal and mitochondrial genes from *Wolbachia* (*16S* and *cox1*), *D. simulans* (*rpl32* and *cox1*), as well as against all the RNA viruses identified upon the annotation analyses. Mapping was performed using BBMap v.37.98 (available at https://sourceforge.net/projects/bbmap/).

### Sequence alignment and phylogenetic analysis

RNA viral sequences identified in *D. simulans* were compared with homologous reference sequences retrieved from the NCBI GenBank database and aligned with MAFF v7.450 (E-INS-I algorithm) [42]. Phylogenetic trees on these data were then inferred using sequences of the conserved RNA-dependent RNA polymerase (RdRp) gene. To this end, both the best-fit model of amino acid substitution and phylogenetic relationships were estimated using the Maximum Likelihood (ML) [43] approach implemented in IQ-TREE v1.6.12 [44]. Nodal support was estimated combining the SH-like approximate likelihood ratio test (SH-aLRT) and the Ultrafast Bootstrap Approximation (UFboot) [45]. Redundant contigs with over 99% amino acid similarity were excluded from the phylogenetic analysis.

### Statistical analysis

The assumption of data normality was assessed by visual inspection and using Kolmogorov-Smirnov (K-S) and Shapiro-Wilk’s tests. As the data was not normally distributed, a Mann-Whitney-Wilcoxon test was used to compare the RNA virome composition with respect to the presence/absence of *Wolbachia*. Comparisons were made using raw and transformed data corresponding to RPM values (i.e. viral abundance) for each library. All the analyses were performed using R software package rstatix.

## Results

A total of 272 female flies were wild-caught in the Perth Hills in Western Australia and tested for *Wolbachia* infection through their F1s. The overall prevalence of *Wolbachia* was 63.6% (173/272), with frequencies at the three sampled locations varying from 54.8% (Raeburn Orchard, N = 73) to 63.8% (Roleystone, N = 130) and 72.5% (Cannington, N = 69). From the Raeburn Orchard field females, we randomly selected a subset of 16 flies representing eight *Wolbachia*-positive and eight Wolbachia-negative specimens for individual sequencing and RNA virus screening.

We identified the *Wolbachia* strain in *D. simulans* using sequence-specific primers targeting the *wsp* gene. We further confirmed the occurrence of *Wolbachia* by mapping the reads back to the *w*Ri and *w*Au *wsp* gene. Most of the *Wolbachia*-infected flies showed a median coverage >100 reads, number of mapping reads >40, and coverage percentage >90% to the reference *w*Au strain, confirming that infected flies harbor *w*Au rather than the wRi strain of *Wolbachia*. No reads mapped to the *wsp* gene for library RAPP88 (**Table S1**) despite the positive infection status determined using a *Wolbachia* specific qPCR assay.

For the sake of comparison of virus diversity among libraries, we mapped the reads of each library to stably expressed genes - *16S* and *cox1* in *Wolbachia* and *rpl32* and *cox1* in *D. simulans*. This provided an internal control to identify any effect on viral abundance due to potential biases introduced during RNA extraction or library preparation. Although, as expected, there was moderate variation in the abundance values, expression levels of reference maker genes were relatively stable across libraries in both *Wolbachia* and *D. simulans* (**Figure 1**).

**Figure 1.**
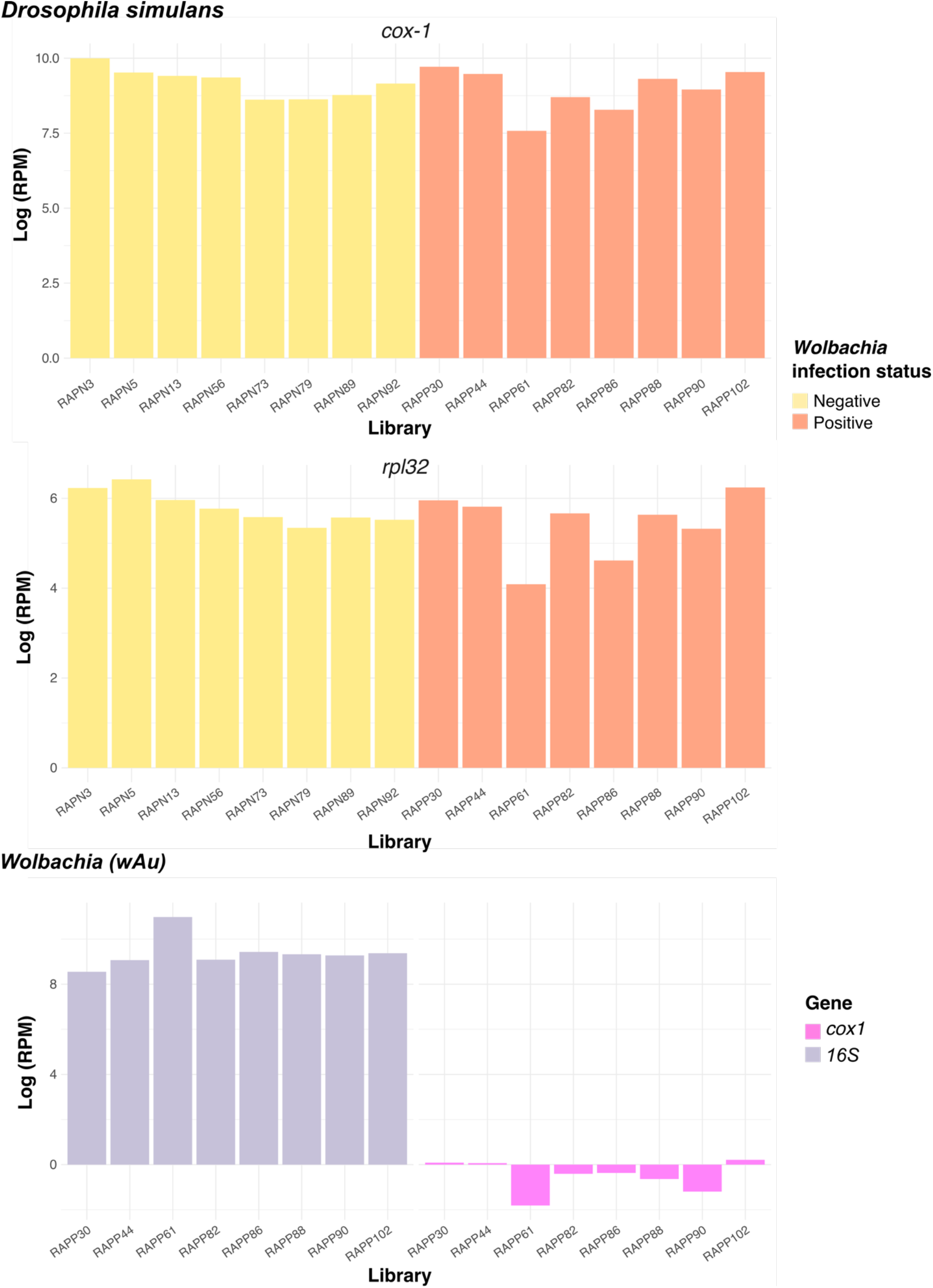
Comparison of the abundance levels of reference genes in *Wolbachia*-positive and *Wolbachia*-negative individual *D. simulans (rpl32 and cox-1)* and *Wolbachia* sp. (*16S* and *cox-1)*.

Overall, we detected ten viruses in the 16 *D. simulans* studied here, five of which were novel (**Figure 2**). Specifically, five viruses shared high sequence identity at the amino acid level (> 96%, e-value = 0.00E+00 - 4.2E-41) to the RdRp of known RNA viruses, whereas the newly discovered viruses shared only between 32.6% to 62.6% amino acid identity to the best viral hit (e-value = 0.00E+00 - 1.4E-06) (**Table 1, Table S4**). Similarly, in five cases phylogenetic analysis of the virus sequences identified revealed close relationships with known *Drosophila*-associated viruses: Galbut virus (*Partitiviridae*), La Jolla virus (*Iflaviridae*), Thika virus (*Picornaviridae*), Nora virus (*Picornaviridae*) and Chaq virus (unclassified) (**Figure 3**). The novel viruses identified, that did not share close phylogenetic relationships to known viruses, were: Raeburn bunya-like virus (*Bunyaviridae*), Araluen mito-like virus (*Mitoviridae*), Carmel mito-like virus (*Mitoviridae*), Lesley reo-like virus (*Reoviridae*), and Cannin tombus-like virus (*Tombusviridae*) (**Figure 3**). Similarity searches against the NCBI/nr database showed that individual flies carried multiple invertebrate-associated viruses from different virus families. For example, up to six viruses were observed in a single *w*Au-negative library (RAPN56) (**Figure 4**, **Table S2**).

**Table 1.**
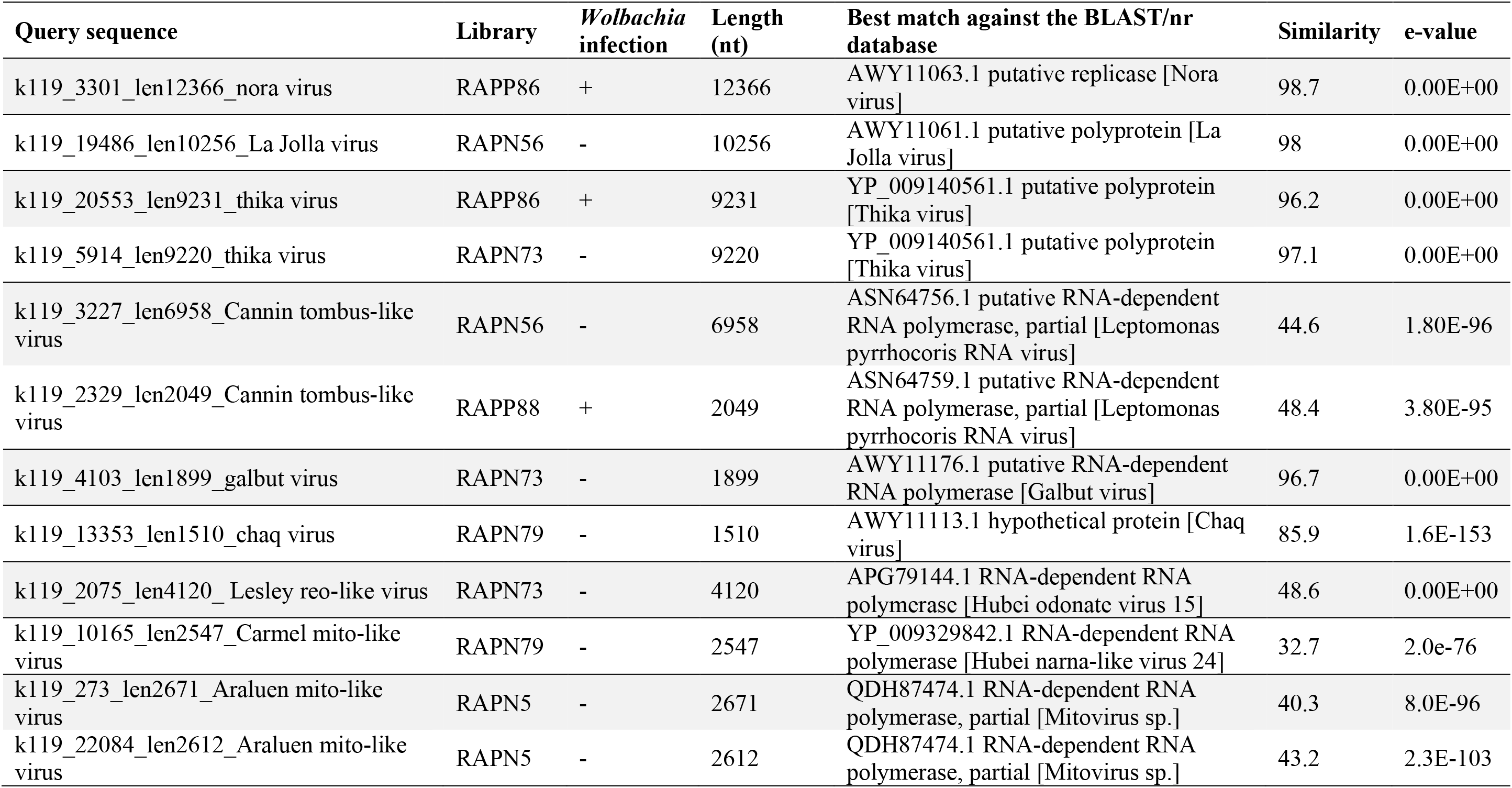
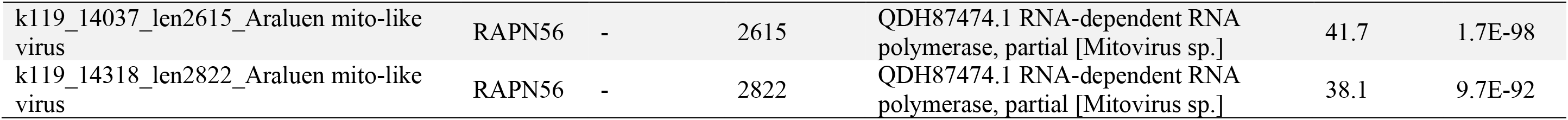
Summary of sequence similarity searches for viruses against the NCBI non-redundant database. Viral sequences listed below correspond to those included in phylogenetic analyses.

**Figure 2.**
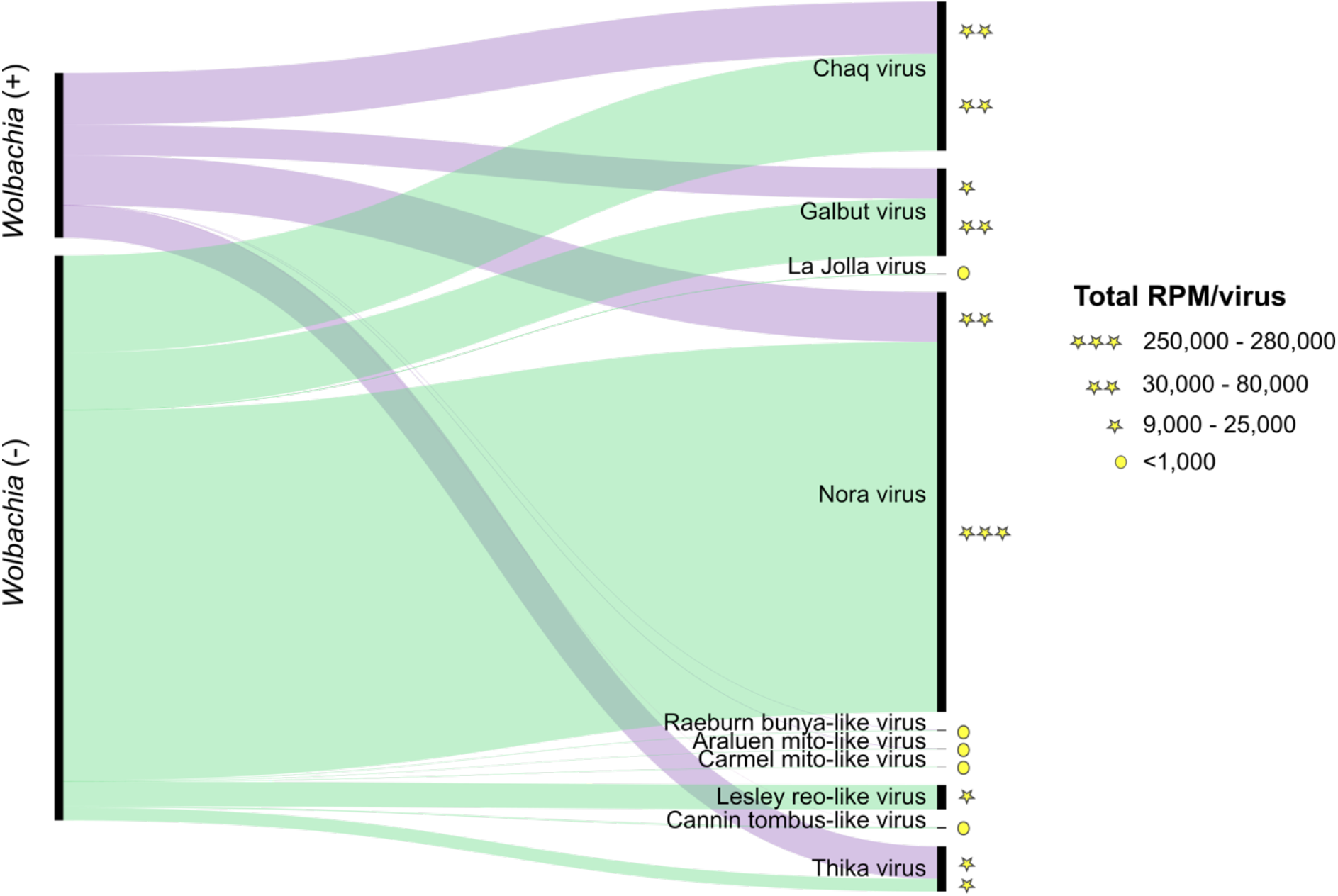
Comparison of viruses found in *Wolbachia*-positive and *Wolbachia*-negative *D. simulans*. The thickness of links is proportional to the total abundance (RPM) of each virus across the samples studied. The range of RPM values are represented with a star and circular shapes.

**Figure 3.**
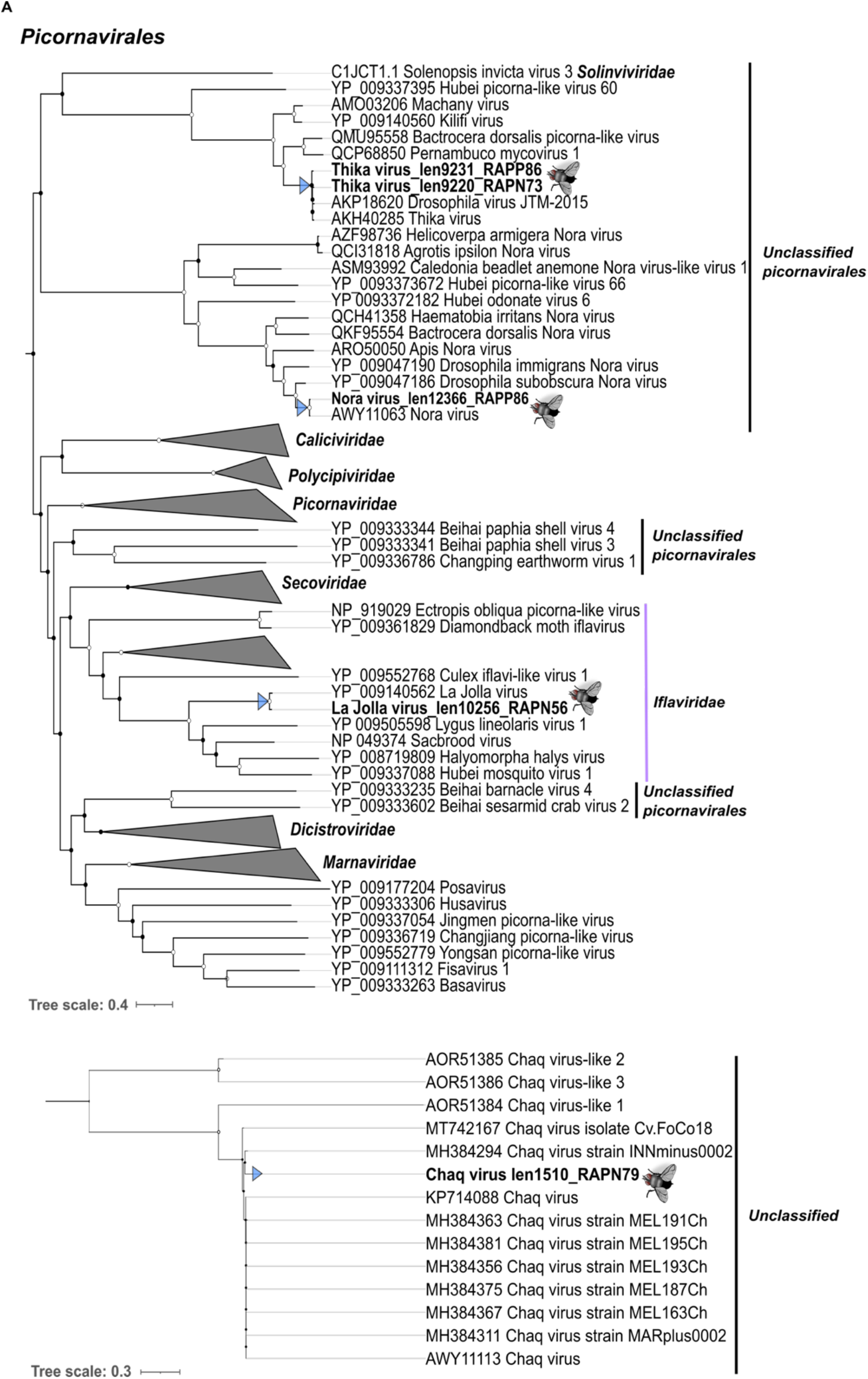
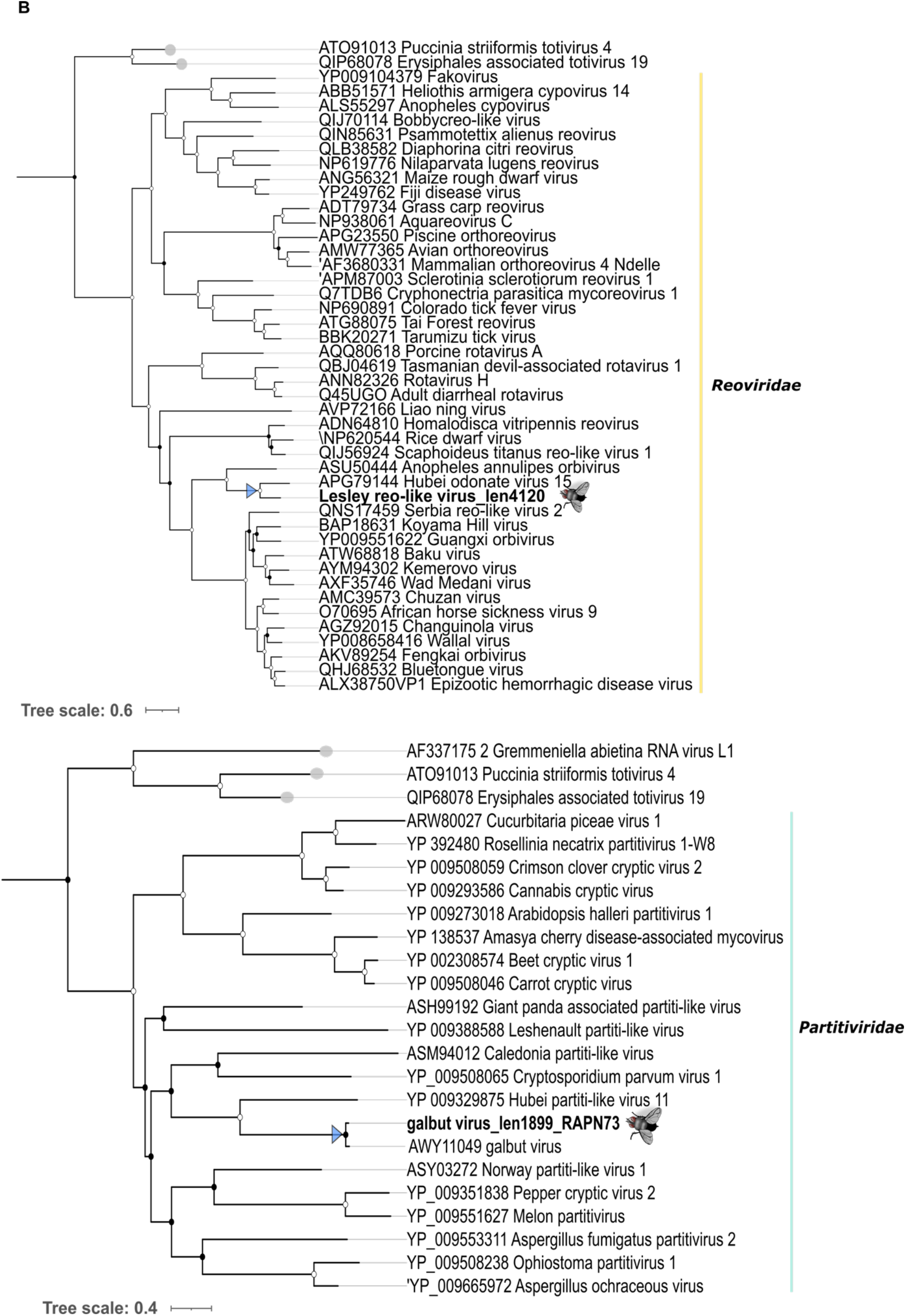
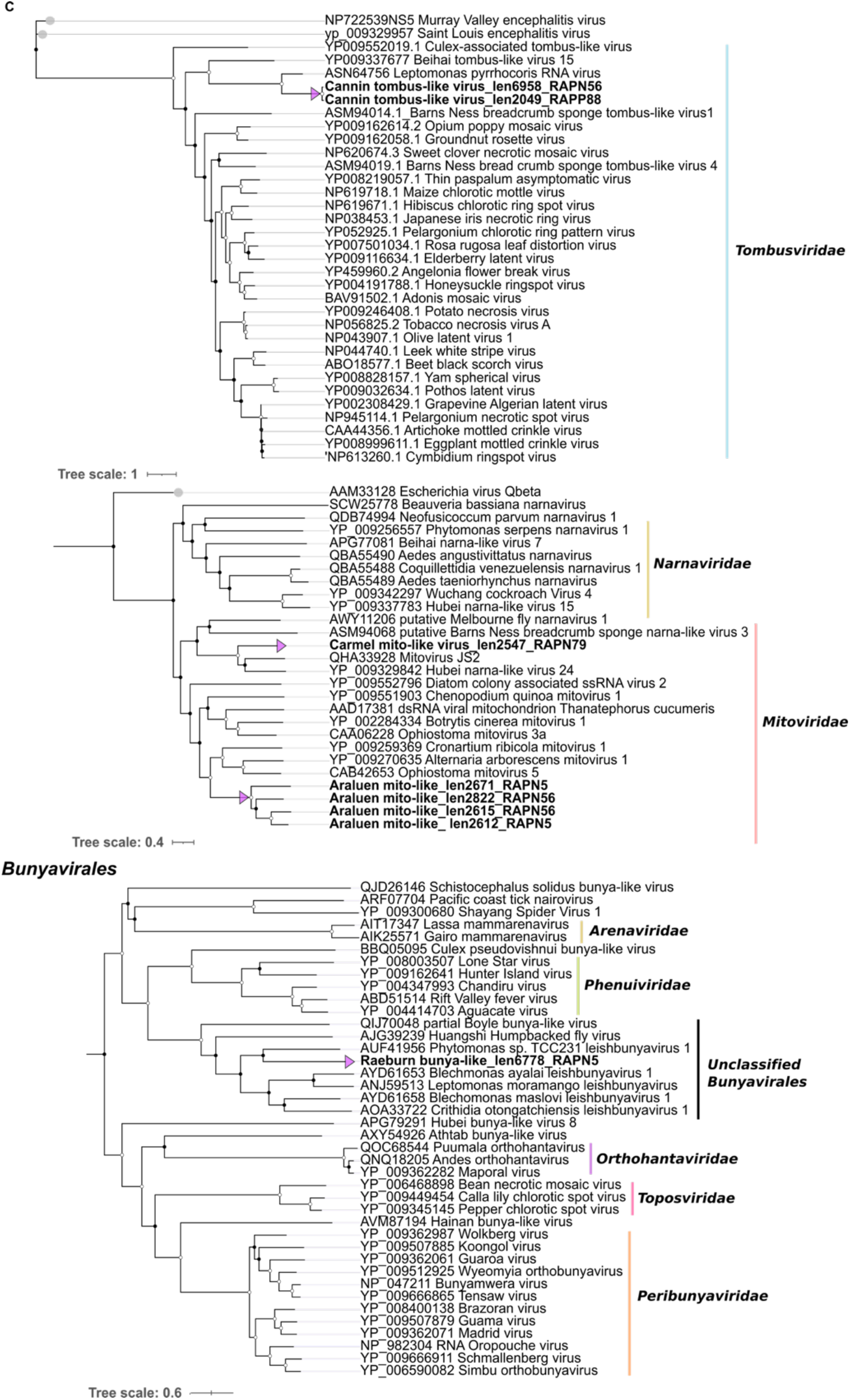
Maximum likelihood phylogenetic trees of the viruses identified from *D. simulans*. The phylogenies were inferred based on the amino acid sequences of the RdRp of seven virus taxonomic groups. Virus family trees were rooted with relevant outgroups that are indicated with grey tips. Order-level trees and the Chaq virus phylogeny (for which no suitable outgroup existed) were midpoint rooted. Coloured arrow tips represent likely (A-B) *Drosophila*-associated viruses and (C) non-*Drosophila*-associated viruses (i.e. that were more likely associated with a component of fly diet or microbiome). Nodal support values greater than 80% (SH-aLRT) and 95% (UFboot) are indicated with white circular shapes at the nodes. Branch lengths are projected using scale bars below each tree.

**Figure 4.**
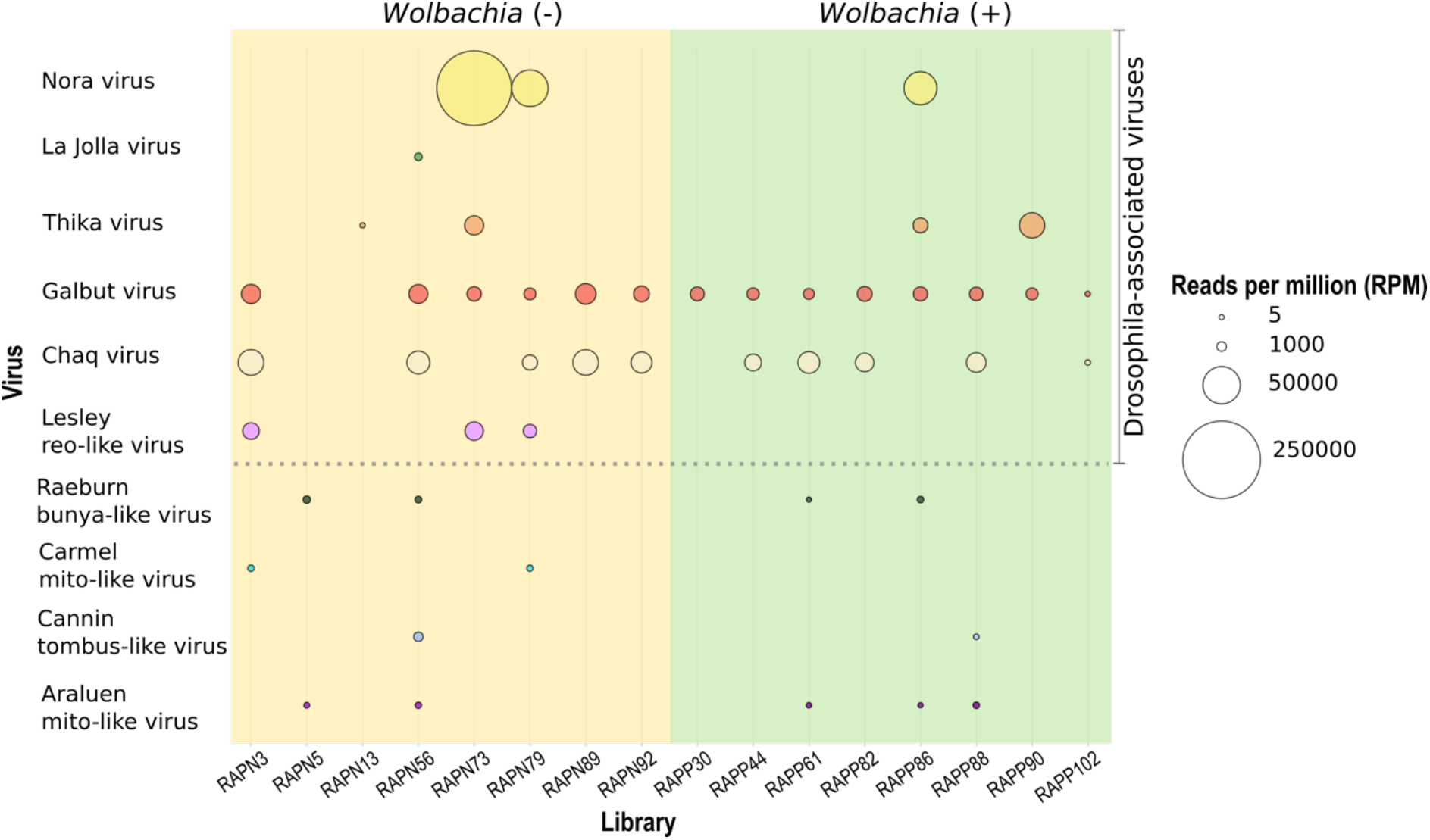
Representation of virome composition and abundance (RPM) across *Wolbachia*-positive and negative libraries. Each library represents an individual *D. simulans* fly. All reads likely due to index-hopping have been excluded.

Some of the newly discovered RNA viruses identified here were likely infecting hosts other than *D. simulans*, and hence might be associated with the fly diet or microbiome. Specifically, these viruses were closely related to Phytomonas sp. TCC231 leishbunyavirus 1 (in the case of Raeburn bunya-like virus), Leptomonas pyrrhocoris RNA virus (Cannin tombus-like virus) and two mito-like viruses (Araluen mito-like virus and Carmel mito-like virus) (**Figure 3, Table S3**), that are associated with trypanosomatid protozoans and fungal hosts, respectively. In contrast, Lesley reo-like virus is likely a *bona fide* arthropod virus. The five newly identified viruses in this study corresponded to full or nearly complete genomes (see below). However, for the majority of the known *Drosophila* viruses we only were able to identify ORFs encoding the RdRp: the exceptions were La Jolla virus and Thika virus for which we also predicted structural components corresponding to coat and capsid proteins.

We next characterized the virome profile present in *D. simulans* in relation to the *w*Au infection status (**Figure 2, Table 1, Table S4**). Accordingly, we identified a slightly higher number (n=10) of viruses in *Wolbachia*-negative flies compared to *Wolbachia*-positive flies (n=7). Among these, Galbut virus, Chaq virus, Nora virus, Thika virus, as well as three novel viruses identified in this study - Raeburn bunya-like virus, Araluen mito-like virus and Cannin tombus-like virus - were present in *D. simulans* regardless of *Wolbachia* infection. In contrast, La Jolla virus, as well as the novel Carmel mito-like virus and Lesley reo-like virus, were only found in wAu-negative flies. Overall, assembled viral contigs displayed high sequence similarity at nucleotide and amino acid level within and between libraries and regardless of the presence/absence of *Wolbachia* (**Table S3**).

We also assessed the potential effect of *Wolbachia* infection on the abundance of RNA viruses present in *w*Au-infected and *w*Au-uninfected flies. Overall, the number of non-rRNA reads represented ~50% of the total of reads (n= 743,389,696 pair-end reads) (**Figure S1**). Furthermore, the RPM values among viruses infecting *Wolbachia* negative and positive infected flies was highly heterogeneous, ranging from 47 to 232,346 and 7 to 37,688 virus RPM, respectively. With the exception of Thika virus, viruses present in both *w*Au-positive and *w*Au-negative flies were 1.87 – 40.17-fold more abundant in the *w*Au-negative individuals than *w*Au-positive *D. simulans.* In contrast, the abundance of Thika virus was 0.39-fold higher in the *Wolbachia*-positive flies (**Figure 3, Table S2**). However, despite this variation in virus abundance levels between groups, there was a non-significant difference between *w*Au-negative and *w*Au-positive *D. simulans* (Mann-Whitney-Wilcoxon test; **Figure 5**). In the case of the viruses only detected in the *w*Au-negative flies, La Jolla virus was present in a single library in moderate abundance (RPM = 378), whilst the newly discovered Lesley reo-like virus was detected in 4/8 libraries (RPM = 3360 - 8749) (**Table S2**).

**Figure 5.**
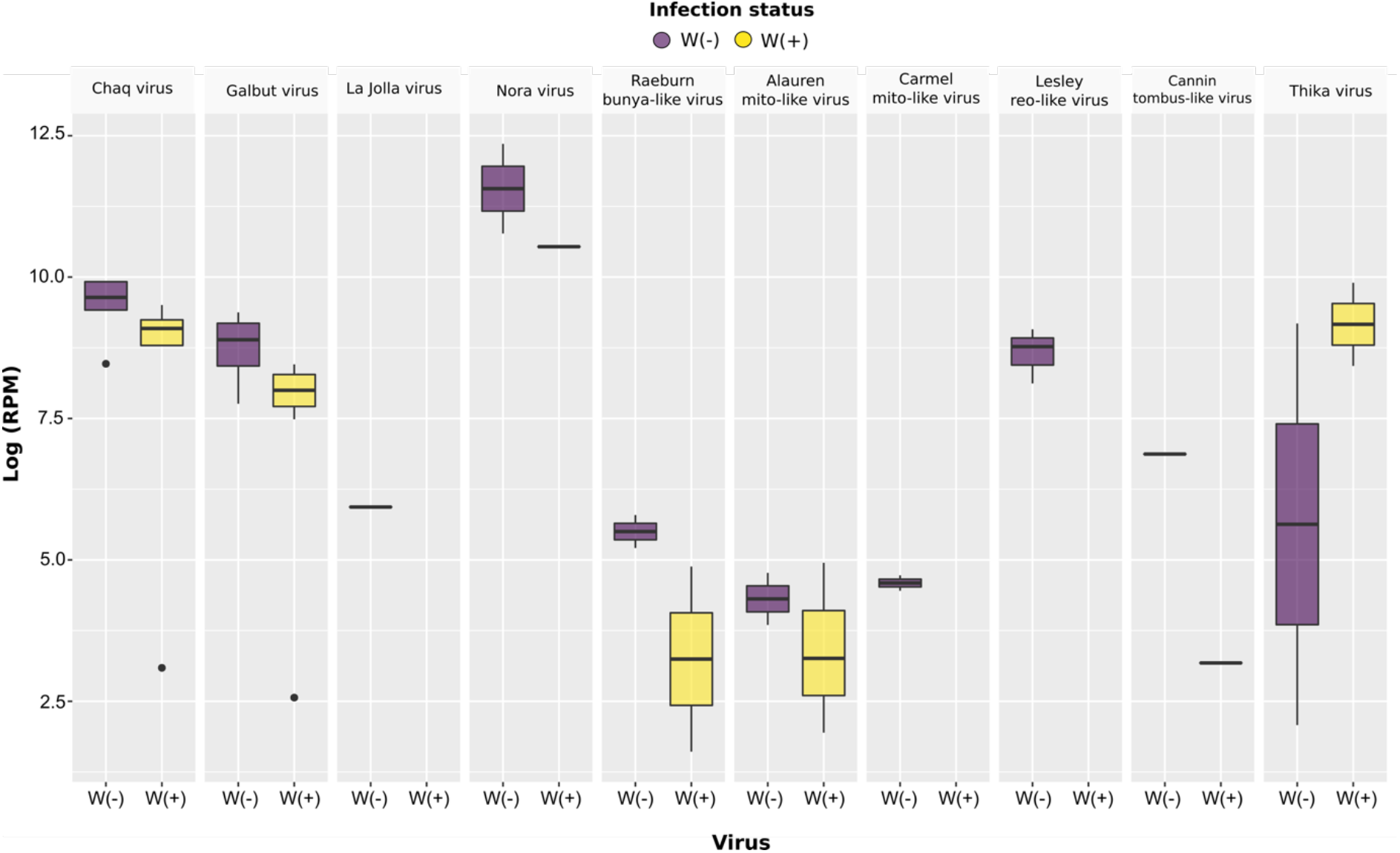
Abundance distribution of seven RNA viruses identified across individual *Wolbachia*-positive and *Wolbachia-*negative *D. simulans*. A non-significant difference was observed between *Wolbachia*-infected and uninfected flies using the Mann-Whitney U test.

## Discussion

The occurrence and spread of *Wolbachia* infection has been widely documented in natural populations of *Drosophila* [10,30,46]. Indeed, *D. simulans* is commonly used as an experimental model to investigate the interactions within the tripartite *Drosophila-Wolbachia*-virus system. In Australia, *D. simulans* can be naturally infected with two *Wolbachia* strains from supergroup A - *w*Au and *w*Ri. While *w*Ri has been gradually displacing *w*Au in eastern Australia, reflected in the changing infection frequencies in surveyed populations since 2004, *D. simulans* from the west coast of Australia only harbor the *w*Au strain [30]. A simple and plausible explanation for this difference is the geographic separation of *D. simulans* populations inhabiting the east and west coasts of Australia and the challenging environmental conditions posed by the intervening desert [30].

We corroborated the presence of *Wolbachia* infection across samples by identifying the *wsp*, *16S* and *cox1* marker genes. The lack of reads mapping to the library RAPP88 might reflect either low levels of *wsp* RNA molecules present in the input for library preparation or high variability compared to the reference sequence. Although *Wolbachia* density was not experimentally assessed, the similar levels of *16S* and *cox-1* abundance across libraries suggest no appreciable biases in the library preparation and RNA sequencing steps.

Estimates from previous surveys showed that the frequency of the *w*Au strain in Western Australia exceeded 50% in *D. simulans* [30]. This is consistent with the data provided here and suggests that *Wolbachia* might be present in a significant proportion of the natural fly population, at least around Perth. Although *w*Au does not cause cytoplasmic incompatibility, its spread is hypothesized to confer fitness advantages (increased survival and/or reproduction) to the host, including antiviral protection [47,48], that might favour its spread and prevent the bacteria from being eliminated from *D. simulans* populations [30,49]. However, our comparison of *Wolbachia*-infected and uninfected *D. simulans* in western Australia revealed no clear effect of *Wolbachia* infection on virome composition and viral abundance between *Wolbachia* infected/uninfected animals. Although our analysis is based on a small sample of individual flies, the apparent absence of a *Wolbachia*-mediated virus protection effect in natural *D. simulans* is compatible with previous findings on *D. melanogaster* naturally infected with *w*Mel in eastern Australia [27], in which virus protection was not observed regardless of the *Wolbachia* infection status and *Wolbachia* density. Even so, the absence of significant association between *w*Au infection and virus diversity does not necessarily translate into a homogeneous effect of *w*Au on the different viruses identified here. For example, it is plausible that the restricted presence of La Jolla virus and the newly identified Lesley reo-like virus in *Wolbachia*-free flies could reflect some impact of antiviral protection in *D. simulans* [27,50]. Indeed, contrasting results were observed in *D. melanogaster*, where La Jolla virus was widely distributed across different libraries [27].

It has previously been shown that the *w*Au strain of *Wolbachia* has a protective role against virus infection in *D. simulans* when flies are challenged with Flock House virus (FHV) and Drosophila C virus (DCV) in a laboratory setting [24,32]. Moreover, the *w*Au strain is protective against the Dengue and Zika viruses in *Aedes aegypti* mosquitoes [51]. Although our observation of an apparent lack of *Wolbachia*-mediated antiviral protection contrasts with those obtained previously, it is likely that differences may depend on *Wolbachia*-host species combinations and natural/artificial viral infections, which may also explain the contrasting results for La Jolla virus. Indeed, most of the available studies have documented the antiviral effect in transinfected insect hosts with non-natural *Wolbachia* strains/viruses under laboratory conditions, as opposed to the study of the natural virome undertaken here. This highlights the importance of careful studies of the interactions within the host-virus-*Wolbachia* system along with environmental factors in natural populations [52–54].

As well as the small sample size, an important caveat of our work is that we explored the *Wolbachia*-mediated virus protection in terms of virus abundance levels reflected in RPM values. This provides insights into virus resistance, but not on tolerance or host survival. Thus, it is still possible that *Wolbachia* is increasing tolerance to virus infection as have been documented for DCV [32]. In addition, although we were not able to assess *Wolbachia* density, previous studies have shown that *w*Au is maintained at high-density in *D. simulans* and has a role on virus blocking [55]. Further research is clearly needed to assess these features in natural populations in order to determine any link with antiviral protection.

Collectively, comparisons of the virome composition in *w*Au infected/uninfected *D. simulans* showed the presence of natural and relatively highly abundant *Drosophila* associated viruses in both groups [25,27,56]. In addition to insect-associated viruses, we identified viruses that are likely to infect other hosts and hence were likely associated with components of *D. simulans* diet or microbiome [57]. For instance, novel viruses from the families *Tombusviridae* and *Bunyaviridae* were related to virus in trypanosomatid protozoa (*Leptomonas* and *Leishmania*). Similarly, given their normal host range distribution, the novel viruses from the family *Narnaviridae* might be associated with fungal hosts. Evidence of trypanosomatids and fungi have been reported in the gut of several species of *Drosophila,* with effects on larvae eclosion and pupation times [57,58]. This, in turn, highlights the extent to which Australian *D. simulans* can be parasitized in nature [58–62].

## Authors and contributors

Conceptualization, M.S., A.A.H. and E.C.H.; methodology, A.S.O.-B., M.S., A.A.H. and E.C.H., formal analysis, A.S.O.-B.; investigation, A.S.O.-B. and M.S.; resources, A.A.H. and E.C.H.; writing—original draft preparation A.S.O.-B.; writing—review and editing A.A.H., M.S and E.C.H.; visualization, A.S.O.-B.; supervision, E.C.H.; All authors have read and agreed to the published version of the manuscript.

## Data summary

The viral genome sequence data generated in this study has been deposited in the NCBI/GenBank database under the accession numbers MW976812-MW976882. Sequence reads are available at the public Sequence Read Archive (SRA) database under the BioProject accession PRJNA706433 (BioSample accessions: SAMN18132282- SAMN18132297).

## Acknowledgements

We thank Vanessa White for research assistance support to conduct this study.

