## Supplementary Information for "RNA virome diversity and *Wolbachia* infection in individual *Drosophila simulans* flies"

**Figure S1**. Ribosomal and non-ribosomal reads in each sequencing library. (A) Number of ribosomal/non-ribosomal reads within the total number of reads. (B) Percentage of ribosomal/non-ribosomal reads within the total number of reads.

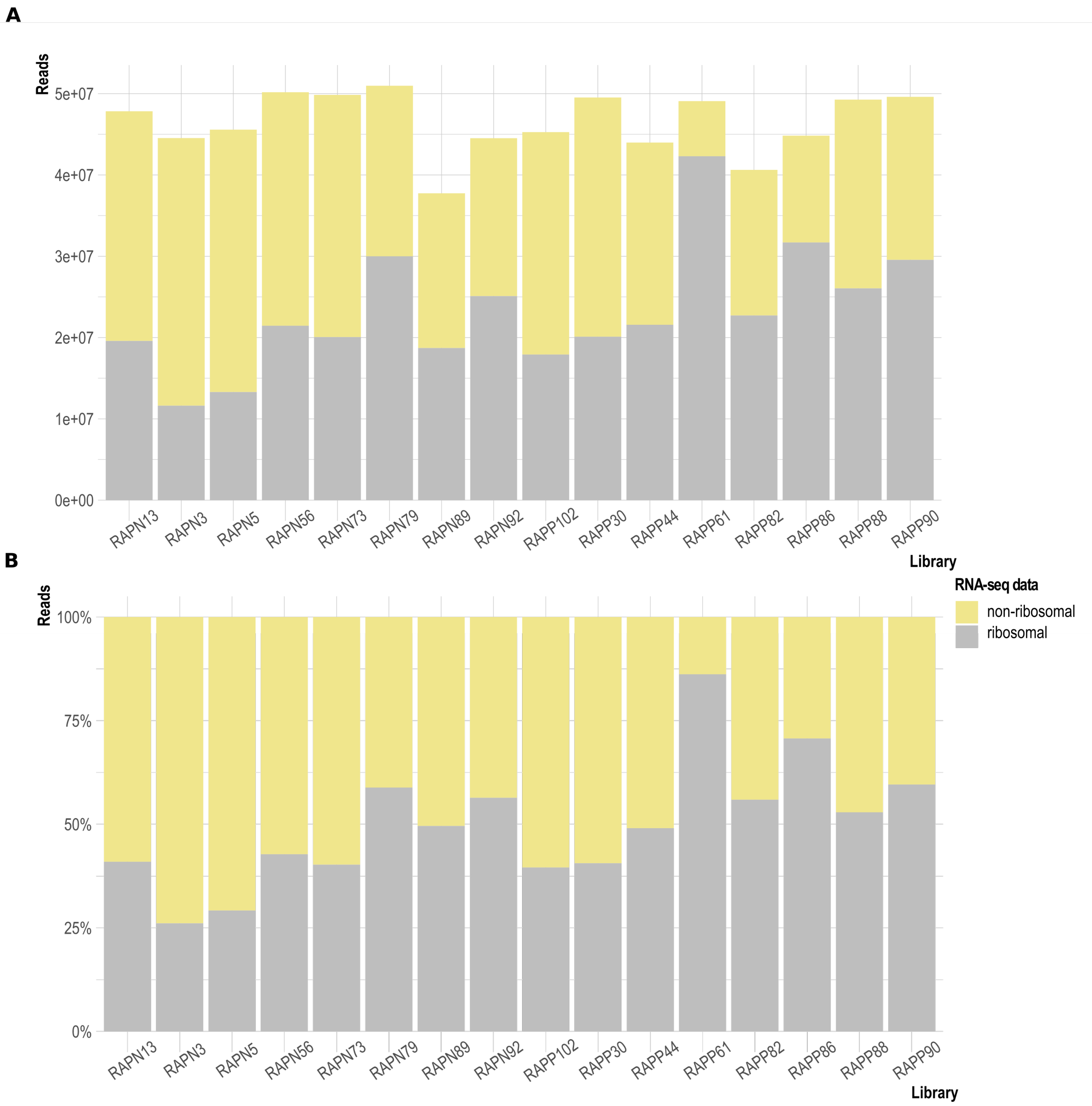

**Table S2.** Abundance values for individual viruses identified in this study across *Wolbachia* infected/uninfected *D. simulans*. The abundance values for a given virus correspond to the number of reads per million mapped reads (RPM). Viruses with RPM values lower than 0.1% of the highest abundance for each virus are shown in grey and assumed to represent index-hopping (and were excluded from additional analyses).

| | ***Wolbachia*-negative *D. simulans*** | | | | | | | |
|---|---|---|---|---|---|---|---|---|
| | **RAPN13** | **RAPN3** | **RAPN5** | **RAPN56** | **RAPN73** | **RAPN79** | **RAPN89** | **RAPN92** |
| **Nora virus (Picorna-like)** | 8* | 5* | 5* | 7* | 232346 | 47585 | 5* | 4* |
| **La Jolla virus (Picorna-like)** | 0 | 0 | 0 | 378 | 0 | 0 | 0 | 0 |
| **Thika virus (Picorna-like)** | 8* | 10* | 14* | 6* | 9694 | 6* | 7* | 5* |
| **Galbut virus (Partiti-like)** | 1* | 9865 | 0 | 9370 | 4264 | 2346 | 11799 | 5650 |
| **Chaq virus (Partiti-like)** | 0 | 20577 | 0 | 15380 | 0 | 4756 | 20293 | 12303 |
| **Lesley reo-like virus** | 0 | 6438 | 0 | 0 | 8749 | 3361 | 1* | 0 |
| **Cannin tombus-like virus** | 0 | 0 | 0 | 964 | 0 | 0 | 0 | 0 |
| **Raeburn bunya-like virus** | 0 | 0 | 328 | 183 | 0 | 0 | 0 | 0 |
| **Carmel mito-like virus** | 0 | 113 | 0 | 0 | 0 | 86 | 0 | 0 |
| **Araluen mito-like virus** | 0 | 0 | 47 | 118 | 0 | 0 | 0 | 0 |
| | ***Wolbachia*-positive *D. simulans*** | | | | | | | |
| | **RAPP102** | **RAPP30** | **RAPP44** | **RAPP61** | **RAPP82** | **RAPP86** | **RAPP88** | **RAPP90** |
| **Nora virus (Picorna-like)** | 1* | 1* | 1* | 0 | 1* | 37688 | 1* | 1* |
| **La Jolla virus (Picorna-like)** | 0 | 0 | 0 | 0 | 0 | 0 | 0 | 0 |
| **Thika virus (Picorna-like)** | 8* | 8* | 7* | 2* | 7* | 4580 | 6* | 19940 |
| **Galbut virus (Partiti-like)** | 13 | 3920 | 2463 | 1784 | 4708 | 3980 | 3597 | 2412 |
| **Chaq virus (Partiti-like)** | 22 | 0 | 6578 | 13454 | 8881 | 15* | 10343 | 0 |
| **Lesley reo-like virus** | 0 | 0 | 0 | 0 | 0 | 0 | 0 | 0 |
| **Cannin tombus-like virus** | 0 | 0 | 0 | 0 | 0 | 0 | 24 | 0 |
| **Raeburn bunya-like virus** | 0 | 0 | 0 | 5 | 0 | 132 | 0 | 0 |
| **Carmel mito-like virus** | 0 | 0 | 0 | 0 | 0 | 0 | 0 | 0 |
| **Araluen mito-like virus** | 0 | 0 | 0 | 26 | 0 | 7 | 141 | 0 |

* RPM values excluded as they likely correspond to index-hopping artefacts during RNA-sequencing.

**Table S3.** List of virus contigs selected for phylogenetic analysis based on levels of sequence similarity.

| Virus | Nº of contigs (Libraries) | % nt identity | % aa identity | Representative sequence (library)* |
|---|---|---|---|---|
| Nora virus (Picorna-like) | 4 (RAPN73, RAPN79, RAPP86) | 99.21 - 100 | 99.65 - 100 | k119_3301_len12366_nora_virus (RAPP86) |
| La Jolla virus (Picorna-like) | 1 (RAPN56) | n/a | n/a | k119_19486_len10256_La_Jolla_virus (RAPN56) |
| Thika virus (Picorna-like) | 3 (RAPN73, RAPP86, RAPP90) | 91.90-99.99 | 96.15 - 99.97 | k119_20553_len9231_thika_virus (RAPP86), k119_5914_len9220_thika_virus (RAPN73) |
| Cannin tombus-like virus | 3 (RAPN56, RAPP88) | 94.34 - 99.27 | 87.44 - 99.41 | k119_2329_len2049_cannin tombus-like virus (RAPP88), k119_3227_len6958_cannin tombus-like virus (RAPN56) |
| Galbut virus (Partiti-like) | 14 (RAPP30, RAPP44, RAPP61, RAPP82, RAPP86, RAPP88, RAPP90, RAPP102, RAPN73) | 96.75-100 | 98.71-100 | k119_4103_len1899_galbut_virus (RAPN73) |
| Chaq virus (Unclassified) | 12 (RAPN3, RAPN56, RAPN56, RAPN79, RAPN89, RAPN92 RAPP102, RAPP44 RAPP61, RAPP82 RAPP86, RAPP88) | 84.48-100 | 98.49-99.69 | k119_13353_len1510_chaq virus (RAPN79) |
| Carmel mito-like virus | 2 (RAPN3, RAPN79) | 98.63 | 99.27 | k119_10165 _len2547_carmel mito-like virus (RAPN79) |
| Araluen mito-like virus | RAPP61, RAPP86, RAPP88, RAPN5, RAPN56 | 39.44-100 | 33.37-98.57 | k119_14037_len2615_araluen mito-like virus (RAPN56), k119_22084_len2612_araluen mito-like virus (RAPN5), k119_14318 _len2822_araluen mito-like virus (RAPN56), k119_273_len2671_araluen mito-like virus (RAPN5) |
| Lesley reo-like virus | 3 (RAPN3, RAPN73, RAPN79) | 99.55 - 99.83 | 88.62 - 99.85 | k119_2075_len4120_lesley reo-like virus (RAPN73) |
| Raeburn bunya-like virus | 3 (RAPN5, RAPN56, RAPP86) | 99.40-99.56 | 99.40-99.54 | k119_6166_len6778_raeburn bunya-like virus (RAPN5) |

**Table S4.** Summary of sequence similarity searches for the total of virus contigs against the NCBI non-redundant database.

| Query sequence | Library | *Wolbachia* infection | Length (nt) | Best match against the BLAST/nr database | Similarity | e-value |
|---|---|---|---|---|---|---|
| k119_18666_len12382_nora virus | RAPN79 | - | 12382 | AWY11063.1 putative replicase [Nora virus] | 98.7 | 0.00E+00 |
| k119_2699_len5328_nora virus | RAPN73 | - | 5328 | AKH67631.1 replication polyprotein [Nora virus] | 98.9 | 0.00E+00 |
| k119_3301_len12366_nora virus | RAPP86 | + | 12366 | AWY11063.1 putative replicase [Nora virus] | 98.7 | 0.00E+00 |
| k119_7728_len7167_nora virus | RAPN73 | - | 7167 | AWY11063.1 putative replicase [Nora virus] | 98.6 | 0.00E+00 |
| k119_19486_len10256_La Jolla virus | RAPN56 | - | 10256 | AWY11061.1 putative polyprotein [La Jolla virus] | 98 | 0.00E+00 |
| k119_5105_len9362_thika virus | RAPP90 | + | 9362 | YP_009140561.1 putative polyprotein [Thika virus] | 96.2 | 0.00E+00 |
| k119_20553_len9231_thika virus | RAPP86 | + | 9231 | YP_009140561.1 putative polyprotein [Thika virus] | 96.2 | 0.00E+00 |
| k119_5914_len9220_thika virus | RAPN73 | - | 9220 | YP_009140561.1 putative polyprotein [Thika virus] | 97.1 | 0.00E+00 |
| k119_6595_len6974_Cannin tombus-like virus | RAPN56 | - | 6974 | ASN64759.1 putative RNA-dependent RNA polymerase, partial [Leptomonas pyrrhocoris RNA virus] | 48.4 | 1.30E-94 |
| k119_3227_len6958_Cannin tombus-like virus | RAPN56 | - | 6958 | ASN64756.1 putative RNA-dependent RNA polymerase, partial [Leptomonas pyrrhocoris RNA virus] | 44.6 | 1.80E-96 |
| k119_2329_len2049_Cannin tombus-like virus | RAPP88 | + | 2049 | ASN64759.1 putative RNA-dependent RNA polymerase, partial [Leptomonas pyrrhocoris RNA virus] | 48.4 | 3.80E-95 |
| k119_14665_len1835_galbut virus | RAPP30 | + | 1835 | AWY11176.1 putative RNA-dependent RNA polymerase [Galbut virus] | 97 | 0.00E+00 |
| k119_7000_len1801_galbut virus | RAPN79 | - | 1801 | AWY11176.1 putative RNA-dependent RNA polymerase [Galbut virus] | 96.1 | 0.00E+00 |
| k119_15592_len2183_galbut virus | RAPN56 | - | 2183 | AWY11176.1 putative RNA-dependent RNA polymerase [Galbut virus] | 96.1 | 0.00E+00 |
| k119_17720_len1823_galbut virus | RAPP88 | + | 1823 | AWY11176.1 putative RNA-dependent RNA polymerase [Galbut virus] | 96.1 | 0.00E+00 |
| k119_4103_len1899_galbut virus | RAPN73 | - | 1899 | AWY11176.1 putative RNA-dependent RNA polymerase [Galbut virus] | 96.7 | 0.00E+00 |

| | | | | | | |
|---|---|---|---|---|---|---|
| k119_17804_len1790_galbut virus | RAPN89 | - | 1790 | AWY11176.1 putative RNA-dependent RNA polymerase [Galbut virus] | 96.1 | 0.00E+00 |
| k119_21494_len1682_galbut virus | RAPP86 | + | 1682 | AWY11176.1 putative RNA-dependent RNA polymerase [Galbut virus] | 96.9 | 0.00E+00 |
| k119_18395_len1758_galbut virus | RAPP82 | + | 1758 | AWY11176.1 putative RNA-dependent RNA polymerase [Galbut virus] | 96.3 | 0.00E+00 |
| k119_15212_len1735_galbut virus | RAPP61 | + | 1735 | AWY11176.1 putative RNA-dependent RNA polymerase [Galbut virus] | 96.1 | 0.00E+00 |
| k119_16819_len1733_galbut virus | RAPP44 | + | 1733 | AWY11176.1 putative RNA-dependent RNA polymerase [Galbut virus] | 95.9 | 0.00E+00 |
| k119_15935_len1648_galbut virus | RAPP102 | + | 1648 | AWY11176.1 putative RNA-dependent RNA polymerase [Galbut virus] | 96.3 | 0.00E+00 |
| k119_3037_len1672_galbut virus | RAPP90 | + | 1672 | AWY11176.1 putative RNA-dependent RNA polymerase [Galbut virus] | 96.1 | 0.00E+00 |
| k119_2472_len1724_galbut virus | RAPN3 | - | 1724 | AWY11176.1 putative RNA-dependent RNA polymerase [Galbut virus] | 96.1 | 0.00E+00 |
| k119_14538_len1827_galbut virus | RAPN92 | - | 1827 | AWY11176.1 putative RNA-dependent RNA polymerase [Galbut virus] | 96.3 | 0.00E+00 |
| k119_21572_len1547_chaq virus | RAPN3 | - | 1547 | AWY11113.1 hypothetical protein [Chaq virus] | 86.2 | 3.7E-154 |
| k119_1063_len1169_chaq virus | RAPN56 | - | 1169 | AWY11113.1 hypothetical protein [Chaq virus] | 87 | 2.8E-122 |
| k119_6487_len519_chaq virus | RAPN56 | - | 519 | AWY11113.1 hypothetical protein [Chaq virus] | 85.7 | 4.2E-41 |
| k119_13353_len1510_chaq virus | RAPN79 | - | 1510 | AWY11113.1 hypothetical protein [Chaq virus] | 85.9 | 1.6E-153 |
| k119_1562_len1480_chaq virus | RAPN89 | - | 1480 | AWY11113.1 hypothetical protein [Chaq virus] | 85.9 | 5.9E-153 |
| k119_12859_len1545_chaq virus | RAPN92 | - | 1545 | AWY11113.1 hypothetical protein [Chaq virus] | 85.9 | 1.6E-153 |
| k119_15738_len1430_chaq virus | RAPP102 | + | 1430 | AWY11113.1 hypothetical protein [Chaq virus] | 85.6 | 5.7E-153 |
| k119_9647_len1467_chaq virus | RAPP44 | + | 1467 | AWY11113.1 hypothetical protein [Chaq virus] | 85.9 | 9.0E-154 |
| k119_18560_ len1471_chaq virus | RAPP61 | + | 1471 | AWY11113.1 hypothetical protein [Chaq virus] | 86.2 | 3.1E-154 |
| k119_17462_len1474_chaq virus | RAPP82 | + | 1474 | AWY11113.1 hypothetical protein [Chaq virus] | 86.2 | 3.1E-154 |

| | | | | | | |
|---|---|---|---|---|---|---|
| k119_4913_ len1396_chaq virus | RAPP86 | + | 1396 | AWY11113.1 hypothetical protein [Chaq virus] | 85.6 | 2.7E-152 |
| k119_2052_len1639_chaq virus | RAPP88 | + | 1639 | AWY11113.1 hypothetical protein [Chaq virus] | 85.6 | 4.7E-151 |
| k119_13097_len4222_Lesley reo-like virus | RAPN79 | - | 4222 | APG79144.1 RNA-dependent RNA polymerase [Hubei odonate virus 15] | 48.6 | 0.00E+00 |
| k119_6812_len4171_ Lesley reo-like virus | RAPN3 | - | 4171 | APG79144.1 RNA-dependent RNA polymerase [Hubei odonate virus 15] | 48.6 | 0.00E+00 |
| k119_2075_len4120_ Lesley reo-like virus | RAPN73 | - | 4120 | APG79144.1 RNA-dependent RNA polymerase [Hubei odonate virus 15] | 48.6 | 0.00E+00 |
| k119_3302_len2571_Carmel mito-like virus | RAPN3 | - | 2571 | YP_009329842.1 RNA-dependent RNA polymerase [Hubei narna-like virus 24] | 32.6 | 3.80E-76 |
| k119_10165_len2547_Carmel mito-like virus | RAPN79 | - | 2547 | YP_009329842.1 RNA-dependent RNA polymerase [Hubei narna-like virus 24] | 32.7 | 2.0e-76 |
| k119_6166_len6778_Raeburn bunya-like virus | RAPN5 | - | 6778 | AUF41956.1 RNA-dependent RNA polymerase [Phytomonas sp. TCC231 leishbunyavirus 1] | 33.8 | 1.50E-225 |
| k119_10640_len6770_Raeburn bunya-like virus | RAPP86 | + | 6770 | AUF41956.1 RNA-dependent RNA polymerase [Phytomonas sp. TCC231 leishbunyavirus 1] | 33.6 | 3.40E-225 |
| k119_13493_len6778_Raeburn bunya-like virus | RAPN56 | - | 6778 | AUF41956.1 RNA-dependent RNA polymerase [Phytomonas sp. TCC231 leishbunyavirus 1] | 33.8 | 1.50E-225 |
| k119_18643_len616_Raeburn bunya-like virus | RAPP61 | + | 616 | ANJ59510.1 putative RNA dependent RNA polymerase [Leptomonas moramango leishbunyavirus] | 50.8 | 5.6E-41 |
| k119_18154_len397_Araluen mito-like virus | RAPP86 | + | 397 | QDH87577.1 RNA-dependent RNA polymerase [Mitovirus sp.] | 41 | 1.3E-17 |
| k119_18803_len660_Araluen mito-like virus | RAPP86 | + | 660 | QDH87474.1 RNA-dependent RNA polymerase, partial [Mitovirus sp.] | 56.6 | 5.1E-56 |
| k119_2431_len529_Araluen mito-like virus | RAPP61 | + | 529 | QDH87474.1 RNA-dependent RNA polymerase, partial [Mitovirus sp.] | 65.5 | 4.2E-13 |
| k119_4889_len444_Araluen mito-like virus | RAPP61 | + | 444 | QDH89956.1 RNA-dependent RNA polymerase, partial [Mitovirus sp.] | 62.6 | 6.3E-34 |
| k119_7925_len640_Araluen mito-like virus | RAPP61 | + | 640 | QDH87474.1 RNA-dependent RNA polymerase, partial [Mitovirus sp.] | 50.5 | 7.9E-46 |
| k119_11923_len1133_Araluen mito-like virus | RAPP61 | + | 1133 | QDH87474.1 RNA-dependent RNA polymerase, partial [Mitovirus sp.] | 37.7 | 1.7E-35 |

| | | | | | | |
|---|---|---|---|---|---|---|
| k119_910_len407_Araluen mito-like virus | RAPP88 | + | 407 | QDH89786.1 RNA-dependent RNA polymerase [Mitovirus sp.] | 43 | 6.8E-11 |
| k119_1774_len2695_Araluen mito-like virus | RAPP88 | + | 2695 | QDH87474.1 RNA-dependent RNA polymerase, partial [Mitovirus sp.] | 40.1 | 1.2E-95 |
| k119_4605_len1802_Araluen mito-like virus | RAPP88 | + | 1802 | QDH87474.1 RNA-dependent RNA polymerase, partial [Mitovirus sp.] | 43.3 | 1.9E-68 |
| k119_14272_len692_Araluen mito-like virus | RAPP88 | + | 692 | QDH86541.1 RNA-dependent RNA polymerase, partial [Mitovirus sp.] | 43 | 1.4E-11 |
| k119_18473_len904_Araluen mito-like virus | RAPP88 | + | 904 | QDH87474.1 RNA-dependent RNA polymerase, partial [Mitovirus sp.] | 47.1 | 2.4E-64 |
| k119_19278_len2543_Araluen mito-like virus | RAPP88 | + | 2543 | QDH87474.1 RNA-dependent RNA polymerase, partial [Mitovirus sp.] | 43.2 | 1.9E-103 |
| k119_19907_len1133_Araluen mito-like virus | RAPP88 | + | 1133 | QDH87474.1 RNA-dependent RNA polymerase, partial [Mitovirus sp.] | 35.5 | 1.4E-29 |
| k119_273_len2671_Araluen mito-like virus | RAPN5 | - | 2671 | QDH87474.1 RNA-dependent RNA polymerase, partial [Mitovirus sp.] | 40.3 | 8.0E-96 |
| k119_2554_len759_Araluen mito-like virus | RAPN5 | - | 759 | QDH86541.1 RNA-dependent RNA polymerase, partial [Mitovirus sp.] | 46.7 | 8.0E-09 |
| k119_2894_len1507_Araluen mito-like virus | RAPN5 | - | 1507 | QDH87474.1 RNA-dependent RNA polymerase, partial [Mitovirus sp.] | 33.3 | 4.4E-43 |
| k119_5165_len1639_Araluen mito-like virus | RAPN5 | - | 1639 | QDH87474.1 RNA-dependent RNA polymerase, partial [Mitovirus sp.] | 41.15 | 4.2E-79 |
| k119_7808_len326_Araluen mito-like virus | RAPN5 | - | 326 | QDH89786.1 RNA-dependent RNA polymerase [Mitovirus sp.] | 42.9 | 2.4E-10 |
| k119_9428_len371_Araluen mito-like virus | RAPN5 | - | 371 | QDH86541.1 RNA-dependent RNA polymerase, partial [Mitovirus sp.] | 38.9 | 1.4E-06 |
| k119_12924_len1504_Araluen mito-like virus | RAPN5 | - | 1504 | QDH87474.1 RNA-dependent RNA polymerase, partial [Mitovirus sp.] | 46.9 | 8.0E-53 |
| k119_22084_len2612_Araluen mito-like virus | RAPN5 | - | 2612 | QDH87474.1 RNA-dependent RNA polymerase, partial [Mitovirus sp.] | 43.2 | 2.3E-103 |
| k119_495 _len2643_Araluen mito-like virus | RAPN56 | - | 2643 | QDH87474.1 RNA-dependent RNA polymerase, partial [Mitovirus sp.] | 43.4 | 2.7E-104 |
| k119_1684_len2692_Araluen mito-like virus | RAPN56 | - | 2692 | QDH87474.1 RNA-dependent RNA polymerase, partial [Mitovirus sp.] | 40 | 8.1E-96 |
| k119_14037_len2615_Araluen mito-like virus | RAPN56 | - | 2615 | QDH87474.1 RNA-dependent RNA polymerase, partial [Mitovirus sp.] | 41.7 | 1.7E-98 |
| k119_14318_len2822_Araluen mito-like virus | RAPN56 | - | 2822 | QDH87474.1 RNA-dependent RNA polymerase, partial [Mitovirus sp.] | 38.1 | 9.7E-92 |
